# A rapid method to determine the bactericidal activity of compounds against non-replicating *Mycobacterium tuberculosis* at low pH

**DOI:** 10.1101/578195

**Authors:** Julie V. Early, Steven Mullen, Tanya Parish

## Abstract

There is an urgent need for new anti-tubercular agents which can lead to a shortened treatment time by targeting persistent or non-replicating bacilli. In order to assess compound activity against non-replicating *Mycobacterium tuberculosis*, we developed a method to detect the bactericidal activity of novel compounds within 7 days. Our method uses incubation at low pH in order to induce a non-replicating state. We used a strain of *M. tuberculosis* expressing luciferase; we first confirmed the linear relationship between luminescence and viable bacteria (determined by colony forming units) under our assay conditions. We optimized the assay parameters in 96-well plates in order to achieve a reproducible assay. Our final assay used *M. tuberculosis* in phosphate-citrate buffer, pH 4.5 exposed to compounds for 7 days; viable bacteria were determined by luminescence. We recorded the minimum bactericidal concentration at pH 4.5 (MBC_4.5_) representing >2 logs of kill. We confirmed the utility of the assay with control compounds. The ionophores monensin, niclosamide, and carbonyl cyanide 3-chlorophenylhydrazone and the anit-tubercular drugs pretomanid and rifampicin were active, while several other drugs such as isoniazid, ethambutol, and linezolid were not.

## Introduction

Tuberculosis, the most fatal disease caused by an infectious agent, caused 1.3 million deaths in 2018 [1]. Treatment requires multiple antibiotics and takes at least 6 months for drug sensitive strains and can take more than 2 years for drug-resistant strains [2]. The long treatment time is at least partly due to non-replicating populations of the causative agent, *Mycobacterium tuberculosis* [3], that are not targeted by most drugs [4]. The inclusion of pyrazinamide to the treatment regimen reduces the duration of therapy from 9 to 6 months; likely because pyrazinamide is active against non-replicating *M. tuberculosis* [5]. Therefore developing new drugs that are active against non-replicating *M. tuberculosis* with the goal to shorten therapy is of utmost importance.

Several laboratory models have been developed to identify compounds active against nonreplicating *M. tuberculosis*. In the Wayne model, *M. tuberculosis* is cultured in sealed glass tubes with a limited headspace ratio; growth of the organism depletes the oxygen content of the medium within two weeks [6]. Entering this hypoxic condition prevents the bacteria from dividing further and initiates a non-replicating, non-dividing state. A high throughput adaptation of this model, the low oxygen recovery assay (LORA), has been used to screen compounds for activity against non-replicating *M. tuberculosis*. LORA includes an outgrowth period of at least 28 hours in normoxia after exposure to hypoxia for 10 days before bacterial survival is measured [7]. Models with more complex conditions have been used to screen compounds. In the multi-stress model, *M. tuberculosis* is cultured using butyrate as a carbon source at pH 5.0, 1% oxygen, and nitric oxide to induce a non-replicating state. After exposure to compounds for 3 days, bacterial survival is assessed by aerobic outgrowth for 7-10 days [8]. An alternative multi stress model used low oxygen (5%), high CO_2_ (10%), low nutrients (10% Dubos medium), and acidic pH (5.0) followed by outgrowth for 9 days and measuring metabolic activity using Alamar blue [9]. A disadvantage of all of these models, is the inclusion of a recovery or outgrowth phase in which bacteria are replicating, which make the interpretation of compound activity more complicated.

More recently, the use of the streptomycin-dependent 18b strain of *M. tuberculosis* allowed the development of an assay which does not use outgrowth [10]. *M. tuberculosis* is deprived of streptomycin for 2 weeks to induce the non-replicating state, bacteria are exposed to compounds for 7 days, and bacterial survival is measured using resazurin [10]. Luciferase has also been used as a readout of bacterial viability, for example using a strain of *M. tuberculosis* expressing an inducible luciferase [11]. Bacteria are maintained in medium without a carbon source for 5 weeks, exposed to compounds for 5 days, and luciferase activity measured in lysed cells with added luciferase reagent [11].

We were interested in developing a simple assay to determine bactericidal activity against nonreplicating *M. tuberculosis*, which did not require an outgrowth period, and which had an easily measured readout with the minimum of manipulations.

## Materials and Methods

### Culture

*M. tuberculosis* H37Rv was cultured in Middlebrook 7H9 medium containing 10% v/v OADC (oleic acid, albumin, dextrose, catalase) (Becton Dickinson) and 0.05% v/v Tween 80 (7H9-OADC-Tw) plus 20 μg/mL kanamycin (Sigma-Aldrich) at 37°C. A recombinant strain carrying plasmid pMV306hsp+LuxAB+G13+CDE (which constitutively expresses luciferase) was used [12, 13]. Phosphate-citrate buffer (PCB) pH 4.5 was prepared by combining 448 mL of 0.2 M dibasic sodium phosphate (Sigma-Aldrich) with 552 mL of 0.1 M citric acid (Sigma Aldrich). The pH was adjusted to pH 4.5 with 0.2 M dibasic sodium phosphate or 0.1 M citric acid solution at room temperature. Tyloxapol (Sigma-Aldrich) was added to a final concentration of 0.05 % v/v; the solution was filter sterilized through a 0.22 μm filter and stored at RT for up to 1 month. CFUs were enumerated by ten-fold serial dilution and plating for colony counts on Middlebrook 7H10 medium containing 10% v/v OADC; CFUs were counted after 4 weeks incubation at 37°C.

### Compound preparation

Compounds were purchased as powders from Sigma-Aldrich, except bedaquiline (Cellagen Technology), linezolid (Selleck Chemicals) and moxifloxacin (Fisher). Compounds were prepared in DMSO at 10 mM and stored in 70 μL aliquots at −20°C; stock solutions were thawed at RT overnight. Two-fold serial dilution series were prepared in 96-well polypropylene plates (Greiner) using a Biomek 3000 or 4000 (Beckman Coulter). Plates were sealed with DMSO-resistant thermal seals (Phenix) and stored at RT for up to 6 months.

### Assay plate preparation

PCB pH 4.5 was dispensed into columns 1-11 (50 μL) and column 12 (100 μL) of sterile 96-well white flat bottom assay plates (Corning Costar) using a Multidrop Combi (Thermo). Compounds (2.04 μL) were added in columns 2-11 using a Biomek 3000 or 4000; rifampicin or DMSO was added into columns 1 and 12 respectively. The final DMSO concentration was 2%, with a typical starting concentration of 200 μM for test compounds. Assay plates were covered with sterile lids and inoculated within 3 h.

### Medium throughput assay for bacterial survival

*M. tuberculosis* was grown to late log-phase (OD_590_ 0.6-1.0), harvested and resuspended in 10 mL of PCB pH 4.5 to an OD_590_ of 1.0; 50 μL of culture was dispensed into each well of columns 1-11 of the assay plates for a final volume of 100 μL per well. Plates were incubated with lids in sealed bags at 37°C in a humidified incubator for 7 days. Luminescence was measured on a Synergy 2 (Biotek) plate reader with dark adjustment for 3 min at RT using a 1 sec integration time.

### Data analysis and quality control

For each assay plate, the Z’ [14] was calculated, and the threshold value was calculated as 3 X the average signal in column 12 (background control). Quality control criteria were set as follows: average RLU for wells in column 12 < 2500; average RLU for wells A1-D1 (maximum kill) < 5000; average RLU for wells E1-H1 (minimum kill) > 50,000; and Z’ was ≥ 0.5. For each compound, the well with the lowest concentration of compound that had RLU below the threshold value was reported as the MBC_4.5_. Based on the RLU to CFU correlation, the MBC_4.5_ is equivalent to ≥ 2 logs decrease in CFU.

## Results

We wanted to develop a rapid and simple assay for determining the bactericidal activity of compounds against *M. tuberculosis* in 96-well format. We selected low pH as the mechanism for inducing the non-replicating state since *M. tuberculosis* is likely to be exposed to acidic conditions in the host [15] and the observation that pyrazinamide shortens treatment and is active against *M. tuberculosis* under low pH conditions [5]. We had previously used standard methods to determine bacterial viability at low pH using CFUs [16], but this is labor intensive and not amenable to testing large numbers of compounds.

### Luminesence is a good readout for bacterial viability

We compared the use of fluorescence and luminescence as a reporter of bacterial viability. Our aim was to find a reporter with a good dynamic range, but in particular with a low limit of detection. We wanted to find a reporter which could reliably monitor a decrease of >2 logs of viable bacteria. We used *M. tuberculosis* constitutively expressing either a fluorescent reporter (*Ds*Red) [17] or the bacterial *lux* operon [13]. The advantage of the latter is that an exogenous substrate is not required [13].

We determined the limit of detection and signal linearity for both strains as compared to OD_590_. OD itself is not a useful readout, since it has a small dynamic range with a lower limit of detection of 0.01 and an upper limit of 1.0. Bacterial cultures were adjusted to OD_590_ ~1.0 and ten-fold serially dilutions prepared in PCB pH 4.5 in 96-well plates (Figure 1). Both reporters demonstrated good signal and linearity over a wide range. However, the luminescence had a larger range of linearity, with a limit of detection about 2 logs lower than for fluorescence (Figure 1). Viable bacteria were measured from the starting cultures; for both H37Rv:*Ds*Red and H37Rv:Luc an OD_590_ of 1.0 was equivalent to 1.7 × 10^9^ ± 1.1 × 10^9^ CFU/mL; for the wild-type strain an OD_590_ of 1.0 was equivalent to 2.8 × 10^8^ ± 1.1 × 10^8^ CFU/mL.

**Figure 1.**
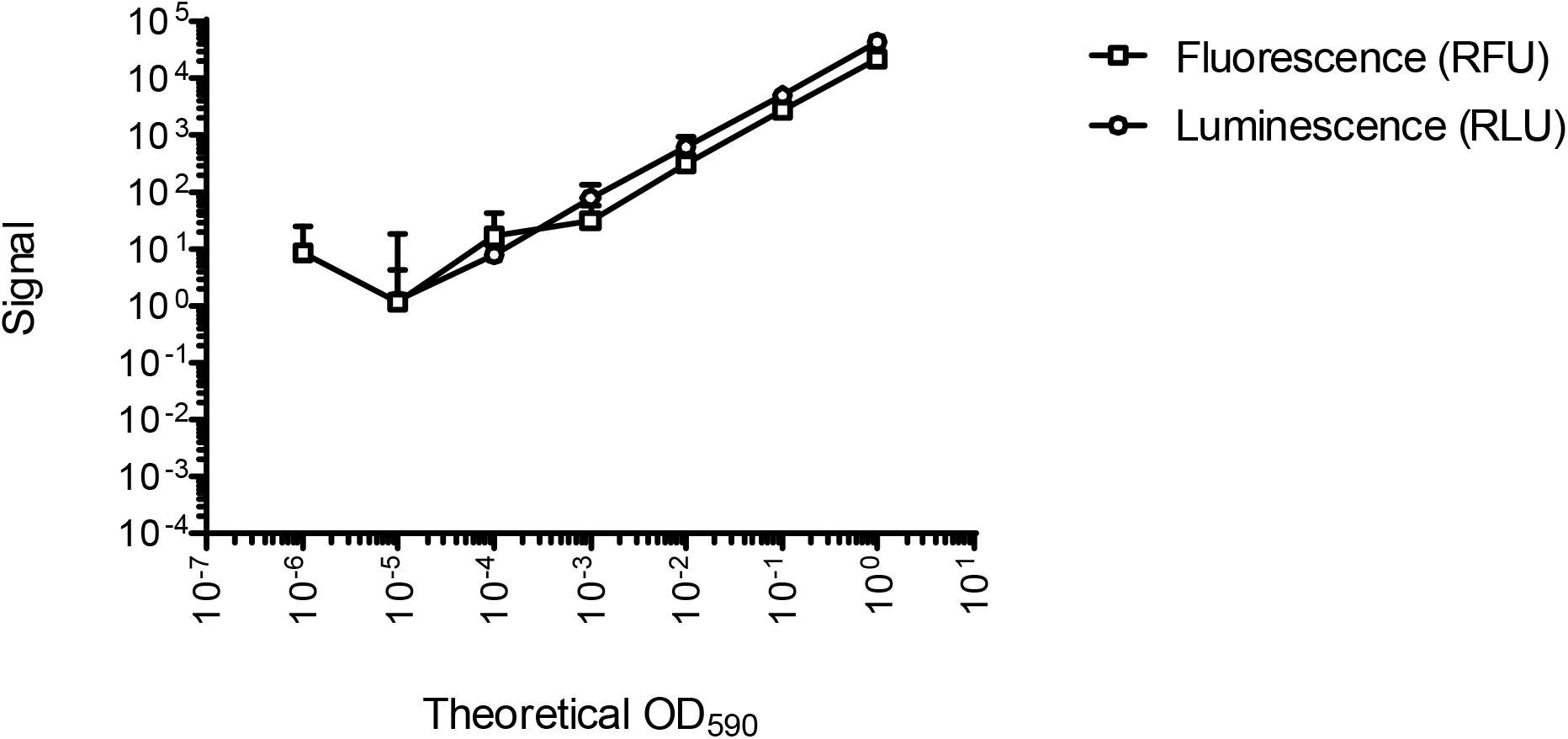
Comparison of fluorescent and luminescent reporters of bacterial viability. Recombinant *M. tuberculosis* strains were grown to late log phase and adjusted to an OD_590_ of 1.0. Serial dilutions were prepared in phosphate citrate buffer in 96-well plates. RLU and RFU were measured using a Synergy H4 plate reader. Data are the average and standard deviation from 2 independent experiments.

We tested the luciferase reporter for its ability to monitor bacterial viability for non-replicating bacteria. We cultured *M. tuberculosis* at low pH over 7 days in PCB pH 4.5 (Figure 2). The limit of detection and linearity did not change after 7 days, with the lower limit at a theoretical OD_590_ of 0.005 and a linear relationship up to an OD_590_ of 0.5. This gave us a usable range of 3 logs difference with a minimum detection of about 8×10^5^ CFU/mL. We saw a good correlation between two independent runs (Figure 2).

**Figure 2.**
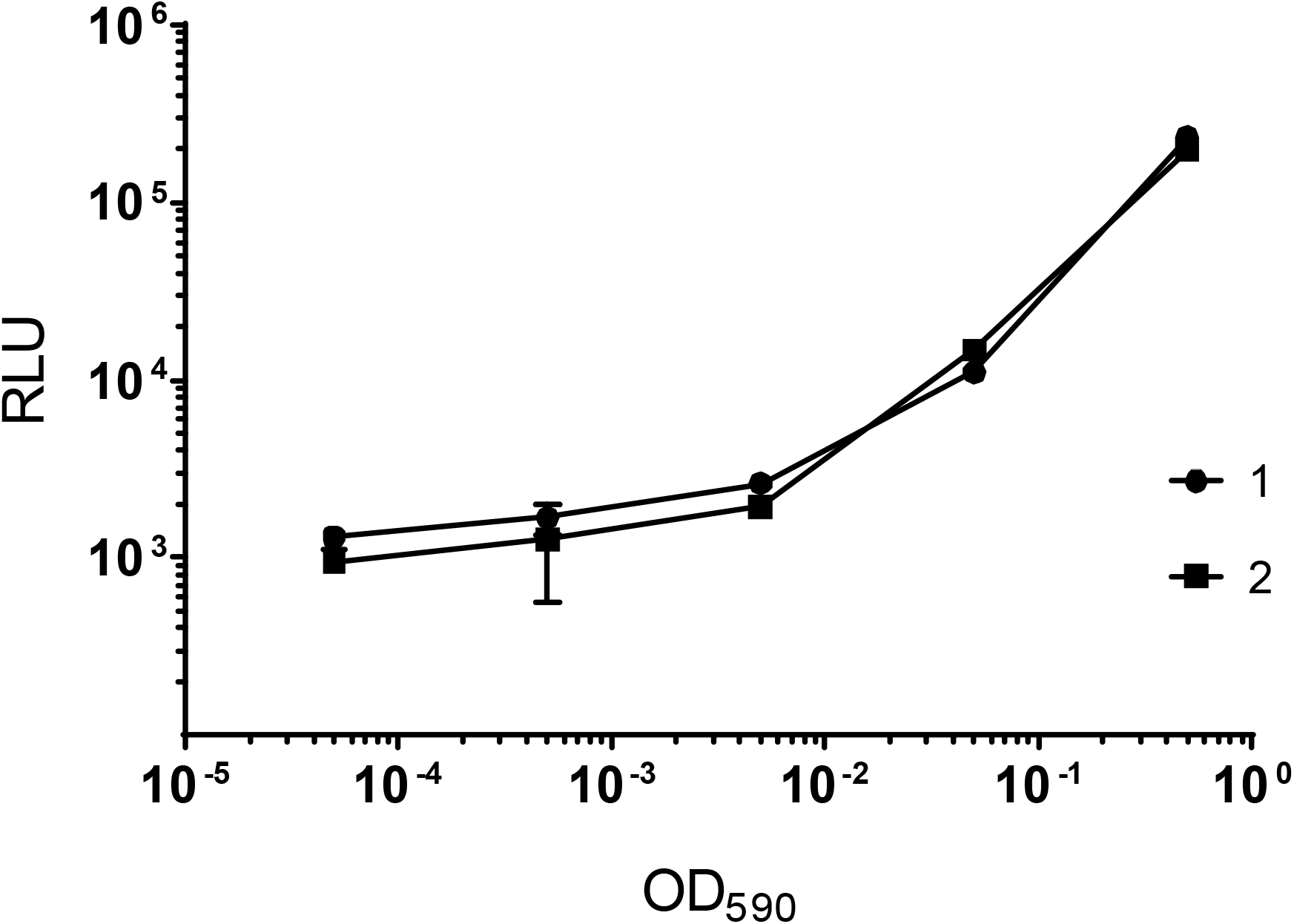
Luminescence as a reporter of bacterial viability in non-replicating *M. tuberculosis*. Recombinant *M. tuberculosis* strains were grown to late log phase and adjusted to an OD_590_ of 1.0. Serial dilutions were prepared in phosphate citrate buffer pH 4.5 and inoculated into 96-well plates. Bacteria were incubated at 37°C for 7 days. RLU was measured using a Synergy 2 plate reader. Data are the average and standard deviation of triplicates from a single experiment. Two independent experiments are shown.

We optimized signal detection using the Synergy 2 plate reader; the final settings were a 3 min dark adjustment, followed by a top read of 1 sec integration. This gave us a signal to background ratio of 104 ± 7.6 (n=3). We monitored assay variability over 21 days. *M. tuberculosis* was incubated in 96-well plates (100 μL per well) for 7-21 days. After 7 days, variability, as measured by the coefficient of variation (CV), was <20% in four independent runs (9%, 10%, 11% and 14%) which meets the criterion for intra-plate variation [18]. Incubation for 14-21 days resulted in higher variation, likely due to increased edge effects from evaporation after prolonged incubation at 37°C (CVs of 23 % at day 14 and 43 % at day 21).

Based on our optimization, we selected the following assay parameters: H37Rv:Luc in PCB pH 4.5 for 7 days of incubation with the Synergy 2 plate reader. The starting OD_590_ was 0.5 with a final assay volume of 100 μL per well and 2% final DMSO concentration using white 96-well plates.

### Luminescence is a reliable reporter of bacterial viability

We tested the ability of the assay to determine bactericidal activity for a range of compounds. We defined bactericidal activity as a 2-log reduction in signal (RLU or CFU) over the 7 days. We set up duplicate plates for each experiment and used one set to measure RLU and the second set to determine CFUs. We tested a total of 156 samples from untreated (DMSO) and treated controls using a range of antibiotics and monensin, an ionophore (Figure 3). For each well, we determined the cutoff point which represents a 2-log reduction in viable bacteria – for RLU we set the cutoff point as 3X the background signal, for CFU it was the experimentally determined numbers. In 87% of the samples, the two readouts gave the same result; there were a small number of discrepancies (13%). Of these, 4 wells (2.6%) were false negatives, where the RLU overestimated the viable bacteria and 17 wells (11%) where the RLU underestimated the viable bacteria. Since the false positive rate was low, we considered that this would be a useful assay to determine bactericidal activity rapidly.

**Figure 3.**
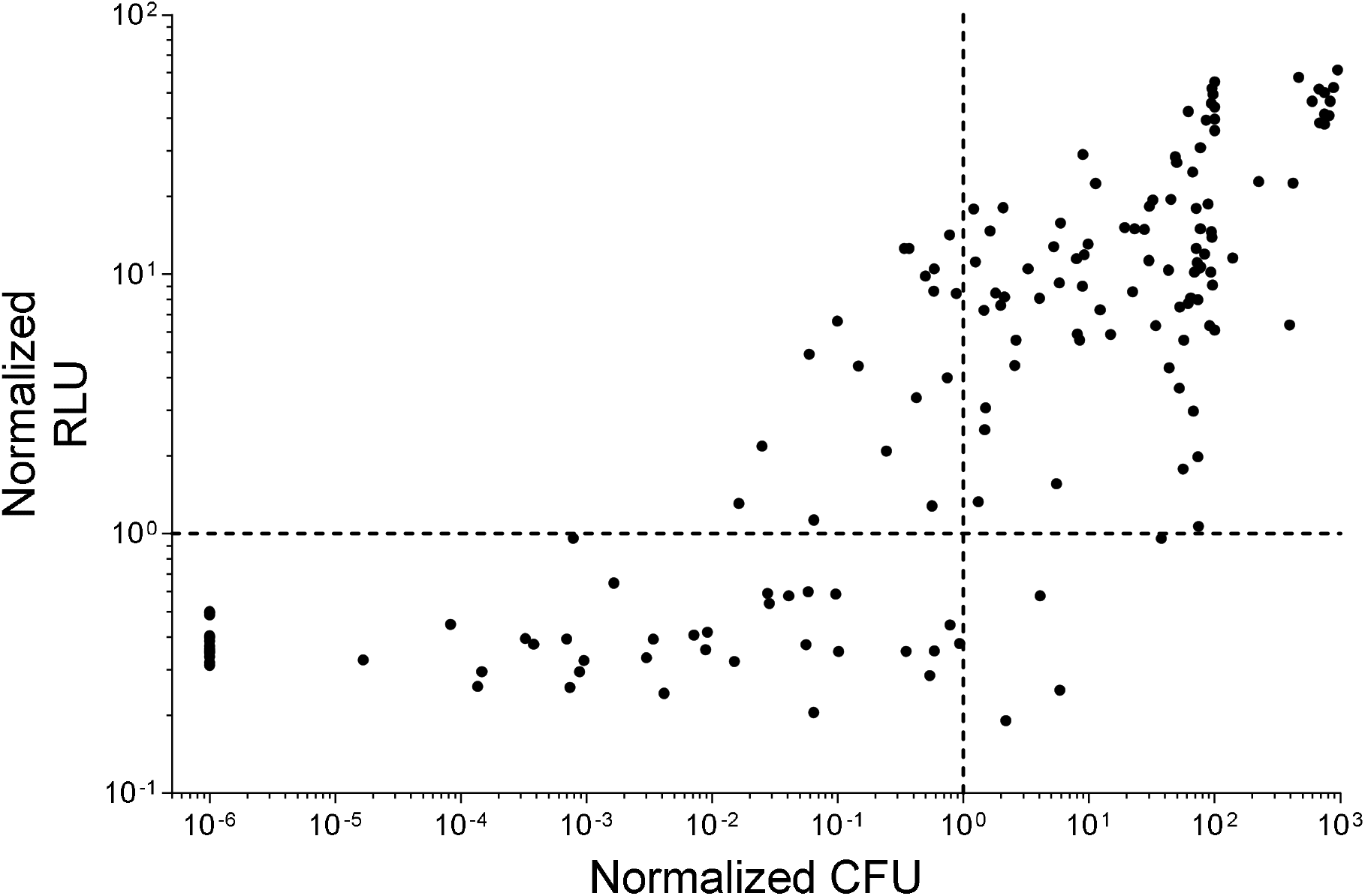
The reliability of luciferase as a marker for viability in non-replicating *M. tuberculosis*. Duplicate assay plates were incubated at 37°C for 7 days. RLU was measured from one set of plates using a Synergy2 plate reader. CFUs were measured from selected wells (n=156). For CFUs, data were normalized to a 2-log reduction from the positive control (DMSO). For RLUs, data were normalized to the threshold level (defined as 3X background from the medium only wells). The dashed lines represent either a 2-log reduction in CFUs or the threshold value for RLUs. The upper left quadrant contains false negatives (11%) and the lower right quadrant contains false positives (2.6%).

### Use of LOPCIDAL to determine MBCs at pH 4.5

We tested a number of known antibiotics and ionophores using this assay, named LOPCIDAL (<u>lo</u>w <u>p</u>H bacteri<u>cidal</u>) (Table 1). Compounds were tested in dose response using two-fold serial dilutions. The MBC at pH 4.5 (MBC_4.5_) was determined as reported as the lowest concentration of compound with a signal below the threshold value. The threshold value was calculated as the average of 3X background for the control column wells (column 12). Data were reproducible between two experiments. This represents a 2-log reduction in viable bacteria. As expected, the ionophores niclosamide, monensin, and CCCP were active with measurable MBC_4.5_s (Table 1). Pretomanid and rifampicin were also active under these conditions, consistent with previous results that these drugs can kill non-replicating *M. tuberculosis* ([19, 20]. Pyrazinamide was not active, despite the fact that we were testing at low pH. However, we were only able to test pyrazinamide up to 200 μM, and higher concentrations of pyrazinamide are required to inhibit growth, even at low pH [21]. Interestingly, the other compounds tested were not active, even known antibiotics and anti-tubercular drugs (isoniazid, bedaquiline and ethambutol). For bedaquiline and metronidazole, this may be due to short length of the assay (7 days) and that these compounds required extended time before killing is observed [22]. These data suggest that killing non-replicating bacilli at low pH may be difficult, and adds further weight to the requirement for new drugs that target bacteria in different physiological states.

**Table 1.** Bactericidal activity of known antibiotics against *M. tuberculosis* at low pH. The MBC_4.5_ was determined for each compound twice; niclosamide and rifampicin were tested multiple times and data reported as average and standard deviation.

| Compound | MBC (μM) | |
| --- | --- | --- |
| Pretomanid | 12 | 6.3 |
| Monensin | 50 | 50 |
| CCCP | 3.1 | 1.6 |
| Bedaquiline | >50 | >50 |
| Metronidazole | >200 | >200 |
| Linezolid | >200 | >200 |
| Pyrazinamide | >200 | >200 |
| Ethambutol | >200 | >200 |
| Levofloxacin | >200 | >200 |
| D-Cycloserine | >200 | >200 |
| Isoniazid | >200 | >200 |
| Moxifloxacin | >200 | >200 |
| Rifampicin | 75 ± 28 (n = 21) |  |
| Niclosamide | 0.22 ± 0.080 (n = 9) |  |

## Conclusion

We have developed a rapid method to identify compounds with bactericidal activity against nonreplicating *M. tuberculosis* at low pH. The assay uses a luciferase reporter as a measure of bacterial viability and can provide data within 7 days. Since the assay uses a 96-well format, it is amenable to automation, can be high throughput, and does not require an outgrowth period. The reduction in time from 49 days (7 weeks) to 7 days is a substantial saving. In addition, the 96-well format allows for full dose response curves to be generated in a less labor intensive fashion. This method could be used to support lead generation programs in drug discovery, where rapid assays for structure-activity relationship studies are required.

Although this assay can be used as a rapid screen, it does not provide full determination of kill kinetics and cannot be used for longer than 7 days (due to edge effects). However, it can be used as a rapid screen to identify molecules that can be followed up in more detail. The main disadvantage of this assay is the higher starting inoculum (>10^8^ CFU/mL), so that compounds with an inoculum-dependent effect might be missed, and that a small population of resistant mutants might be present at the outset [23]. The latter issue is tempered by the fact with the assay only requires 7 days growth, and generally resistance is not seen in outgrowth until day 14. In addition this assay is carried out under non-replicating conditions, such that resistant mutants will not grow.

## Acknowledgements

We thank Torey Alling for technical assistance. This work was funded in part by Eli Lilly and Company in support of the mission of the Lilly TB Drug Discovery Initiative and by the Bill and Melinda Gates Foundation, under grant OPP1024038.

## Author Contributions

JE contributed to methodology, investigation, supervision, formal analysis, validation, visualization, writing the first draft, and editing. SM contributed to methodology, investigation, validation, visualization, analysis, and writing the first draft. TP contributed by conceptualization, formal analysis, validation, providing resources, funding acquisition, supervision, review and editing.

